## Supplementary Figures for "Recovery after human bone marrow mesenchymal stem cells (hBM-MSCs)-derived extracellular vesicles (EVs) treatment in post-MCAO rats requires handling associated with repeated behavioral testing"

**Figure S1. All animals recover their weight loss following MCAO surgery after 28 days post stroke (dps), independently of treatment**

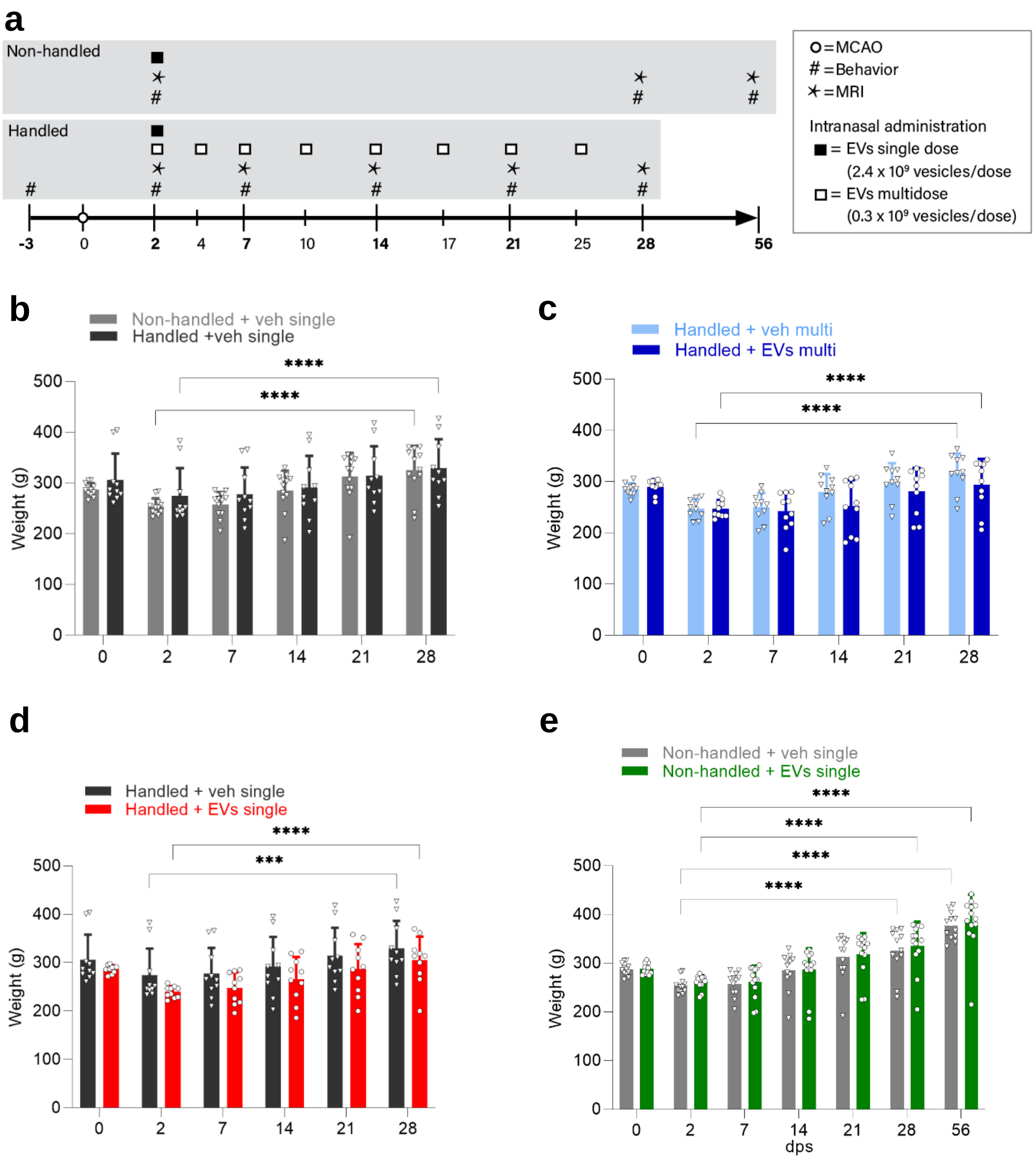

**Figure S1. All animals recover their weight loss following MCAO surgery after 28 days post stroke (dps), independently of treatment.**

**(a)** Experimental design and timeline with the different cohorts and treatments. The exact number of adult MCAO rats used in each cohort and their treatment is specified in **Table S1**.

**(b)** Weights (g) of non-handled and handled rats, receiving a single intranasal dose of vehicle, at 0, 2, 7, 14, 21, and 28 dps.

**(c)** Weights (g) of handled rats, receiving a multidose of vehicle or EVs, at 0, 2, 7, 14, 21, and 28 dps.

**(d)** Weights (g) of handled rats, receiving a single dose of vehicle or EVs, at 0, 2, 7, 14, 21, and 28 dps.

**(e)** Weights (g) of non-handled rats, receiving a single dose of vehicle or EVs, at 0, 2, 7, 14, 21, 28, and 56 dps.

All data was analyzed by two-way repeated measures ANOVA followed by the post hoc Holm-Sidak's multiple comparisons test (***p<0.0005, ****p<0.0001). Bars show mean +/− SD. Each symbol represents a single rat: non-handled + veh single (*n=12*), non-handled + EVs single (*n=13*), handled + veh multi (*n=10*), handled + EVs multi (*n=10*), handled + veh single (*n=10*), handled + EVs single (*n=10*).

**Figure S2. Animals improve spontaneously the total ischemic volume and percentage of edematous expansion of the ipsilateral hemisphere at 28 dps, independently of the treatment.**

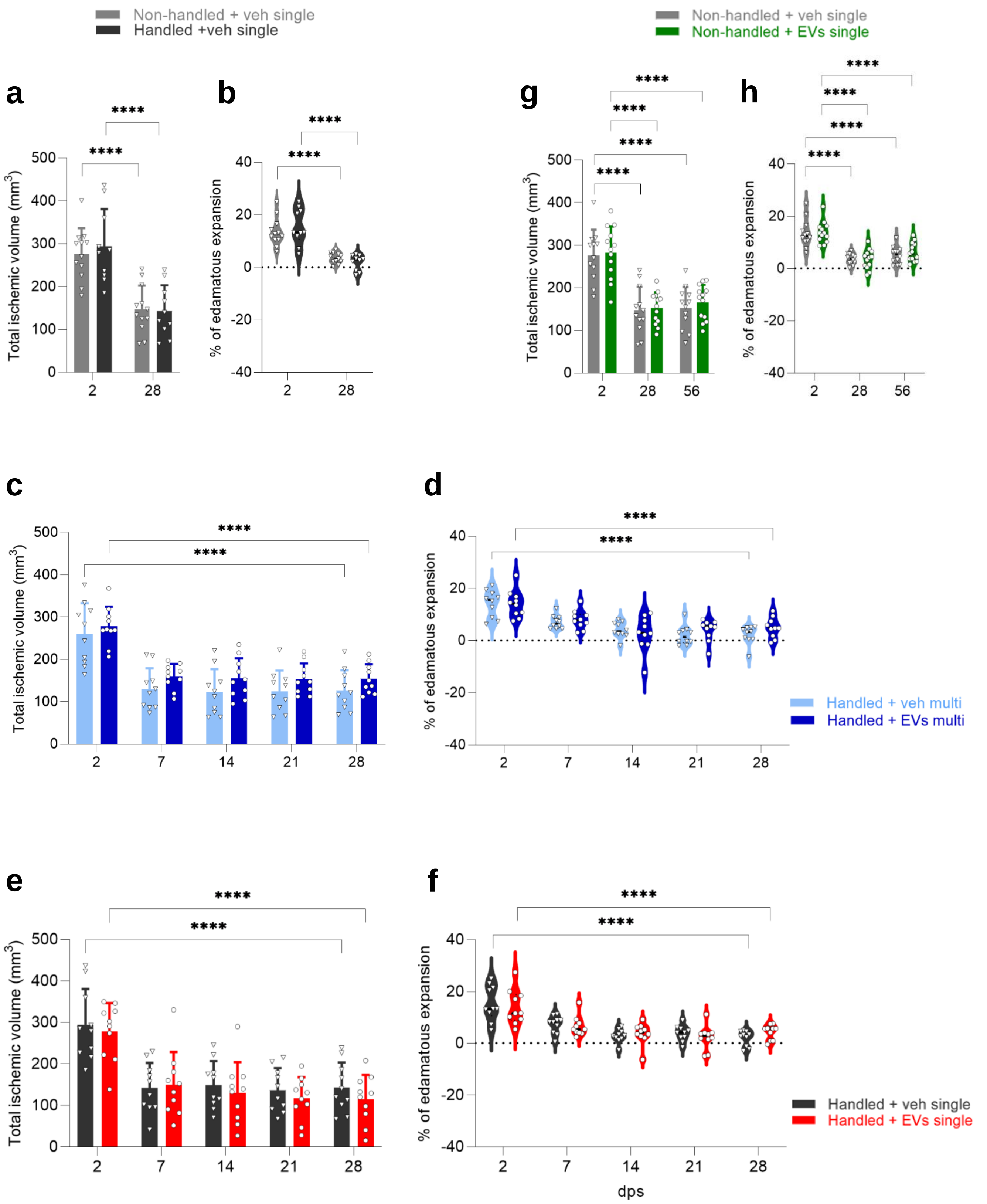

**Figure S2. Animals improve spontaneously the total ischemic volume and percentage of edematous expansion of the ipsilateral hemisphere at 28 dps, independently of the treatment.**

**(a–b)** Total ischemic volume and percentage of edematous expansion of the ipsilateral hemisphere in non-handled and handled rats, receiving a single intranasal dose of vehicle, at 2 and 28 dps.

**(c–d)** Total ischemic volume and percentage of edematous expansion of the ipsilateral hemisphere in handled rats, receiving a multidose of vehicle or EVs, at 2, 7, 14, 21, and 28 dps.

**(e–f)** Total ischemic volume and percentage of edematous expansion of the ipsilateral hemisphere in handled rats, receiving a single dose of vehicle or EVs, at 2, 7, 14, 21, and 28 dps.

**(g–h)** Total ischemic volume and percentage of edematous expansion of the ipsilateral hemisphere in non-handled rats, receiving a single dose of vehicle or EVs, at 2, 28, and 56 dps

All data was analyzed by two-way repeated measures ANOVA followed by the post hoc Holm-Sidak's multiple comparisons test (****$p<0.0001$). Bars show mean +/− SD. Violin plot graphs represent quartiles and median. Each symbol represents a single rat: non-handled + veh single (*n=12*), non-handled + EVs single (*n=13*), handled + veh multi (*n=10*), handled + EVs multi (*n=10*), handled + veh single (*n=10*), handled + EVs single (*n=10*).

### Figure S3. Handled rats treated with multidose EVs do not recover in the cylinder, grid walking, or forelimb placing tests at 28 dps

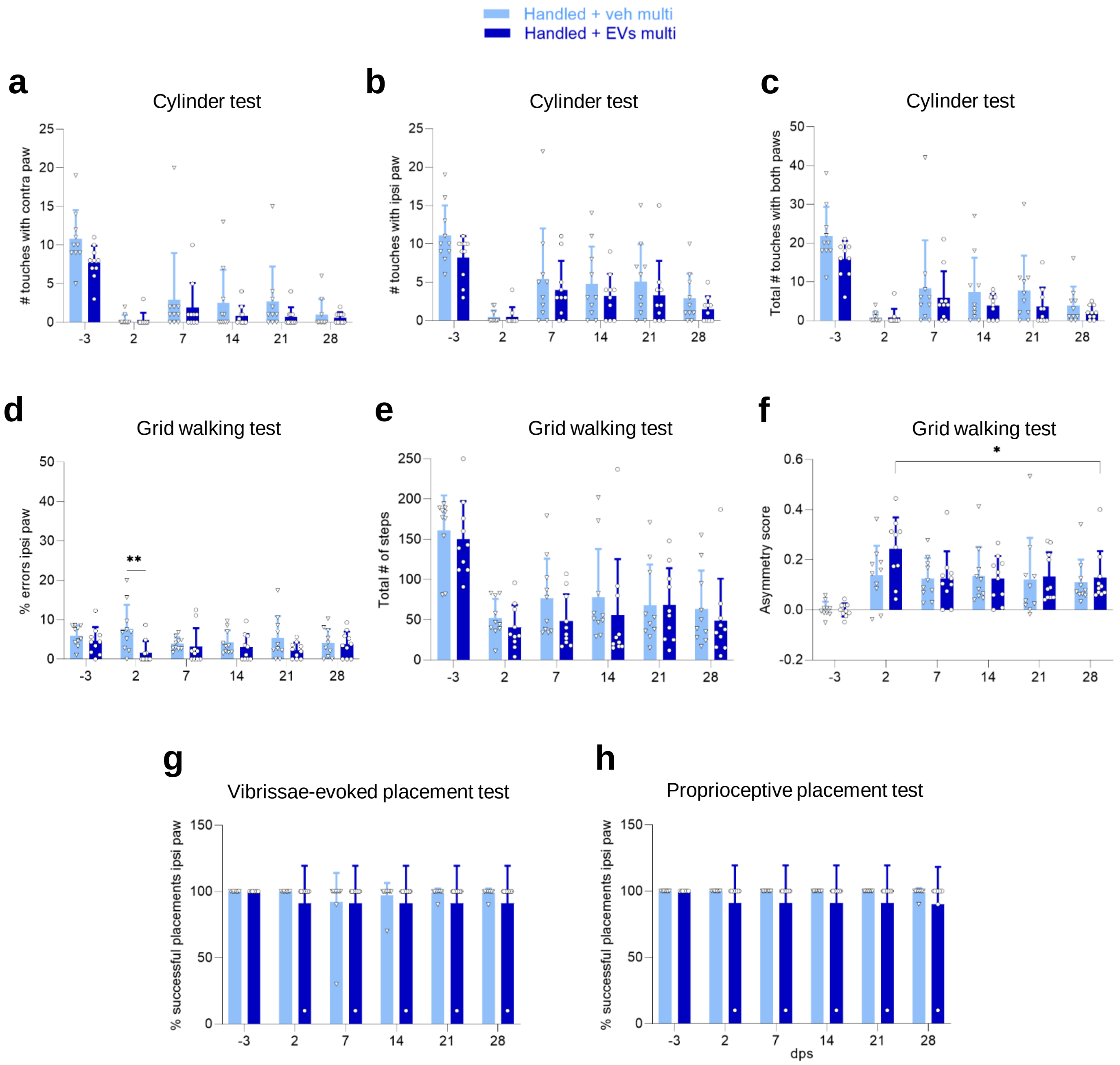

**Figure S3. Handled rats treated with multidose EVs do not recover in the cylinder, grid walking, or forelimb placing tests at 28 dps.**

**(a–h)** Handled rats treated with a multidose of vehicle or EVs were tested at -3, 2, 7, 14, 21, and 28 dps on **(a)** the number of touches with the contralateral paw in the cylinder test, **(b)** the number of touches with the ipsilateral paw in the cylinder test, **(c)** the total number of touches with both paws in the cylinder test, **(d)** the percentage of errors with the ipsilateral foot in the walking grid test, **(e)** the total number of steps in the walking grid test, **(f)** the asymmetry score in the walking grid test, **(g)** the percentage of successful placements with the ipsilateral paw in the vibrissae-evoked forelimb placing test, and **(h)** the percentage of successful placements with the ipsilateral paw in the proprioceptive forelimb placing test.

All data was analyzed by two-way repeated measures ANOVA followed by the post hoc Holm-Sidak's multiple comparisons test (*$p<0.05$, **$p<0.005$). Bars show mean +/− SD. Each symbol represents a single rat: handled + veh multi (*n=10*), handled + EVs multi (*n=10*).

### Figure S4. Handled rats treated with a single dose of EVs do not recover in cylinder, grid walking, or forelimb placing tests at 28 dps

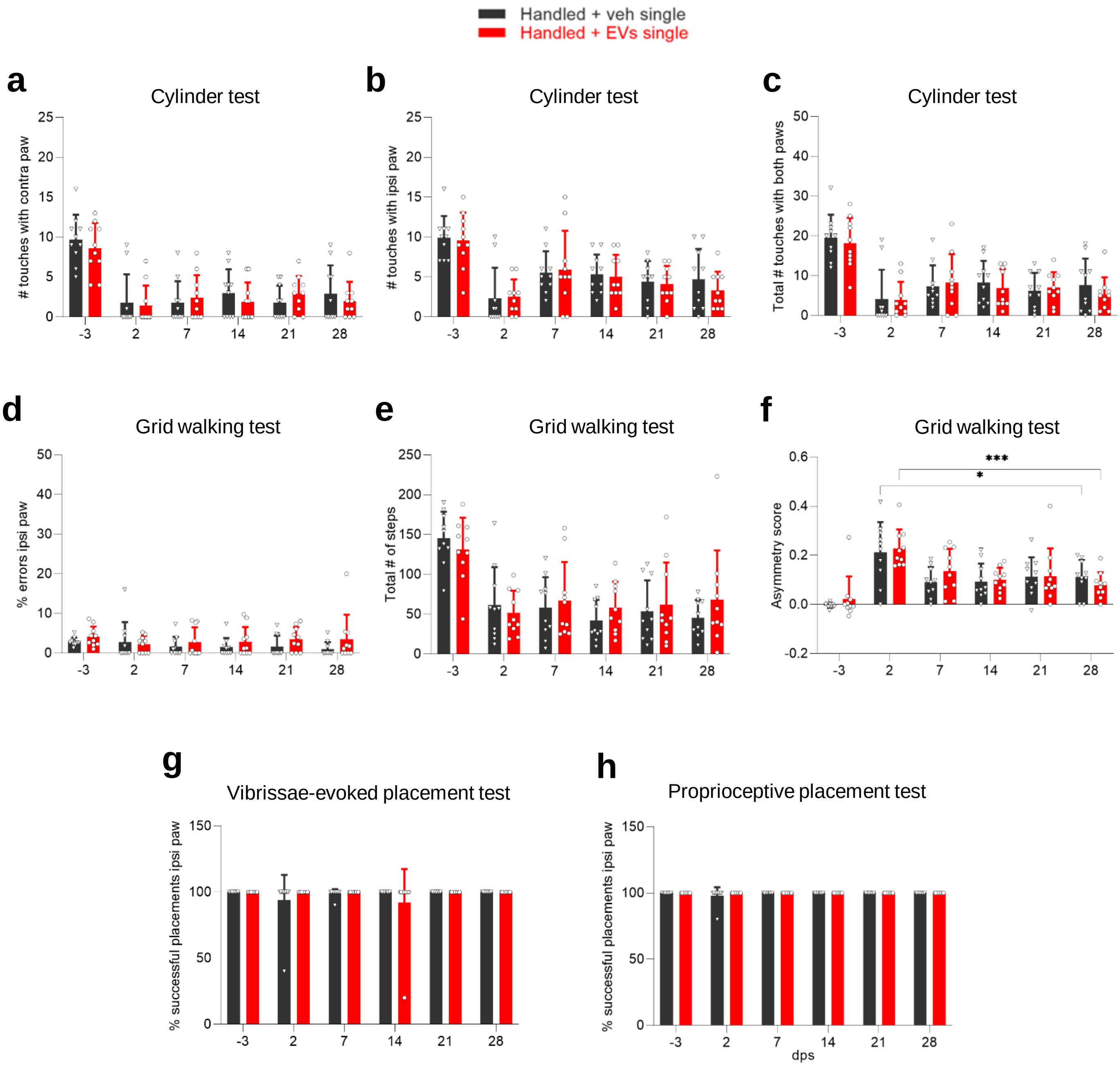

**Figure S4. Handled rats treated with a single dose of EVs do not recover in cylinder, grid walking, or forelimb placing tests at 28 dps.**

**(a–h)** Handled rats treated with a single dose of vehicle or EVs were tested at -3, 2, 7, 14, 21, and 28 dps on **(a)** the number of touches with the contralateral paw in the cylinder test, **(b)** the number of touches with the ipsilateral paw in the cylinder test, **(c)** the total number of touches with both paws in the cylinder test, **(d)** the percentage of errors with the ipsilateral foot in the walking grid test, **(e)** the total number of steps in the walking grid test, **(f)** the asymmetry score in the walking grid test, **(g)** the percentage of successful placements with the ipsilateral paw in the vibrissae-evoked forelimb placing test, and **(h)** the percentage of successful placements with the ipsilateral paw in the proprioceptive forelimb placing test.

All data was analyzed by two-way repeated measures ANOVA followed by the post hoc Holm-Sidak's multiple comparisons test (*p<0.05, ***p<0.0005). Bars show mean +/− SD. Each symbol represents a single rat: handled + veh single (*n=10*), handled + EVs single (*n=10*).

**Figure S5. Handled rats treated with a single dose of EVs do not recover in the mNSS at 28 dps compared to handled rats treated with multidose**

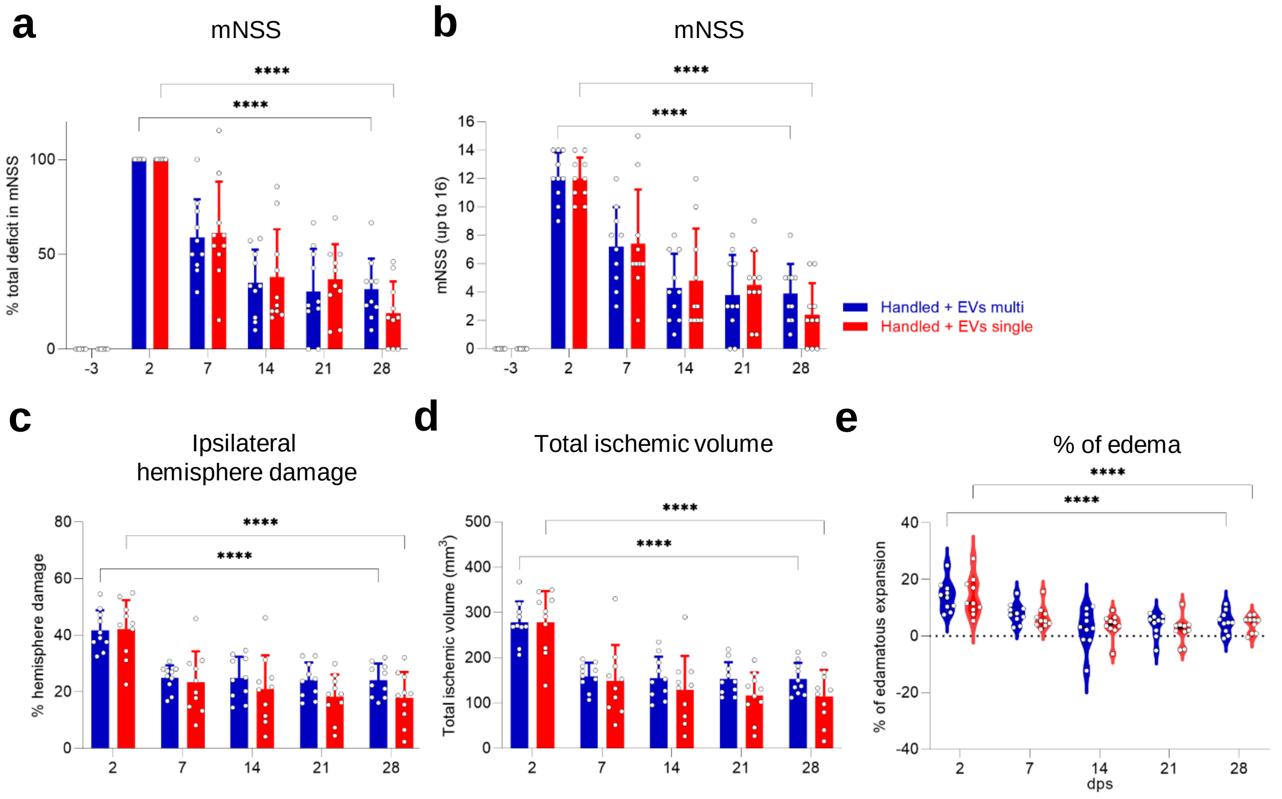

**Figure S5. Handled rats treated with a single dose of EVs do not recover in the mNSS at 28 dps compared to handled rats treated with multidose.**

**(a)** Functional recovery in handled rats receiving a single or multidose of EVs after MCAO stroke, shown as a percentage in total deficit in the modified Neurological Severity Score (mNSS) at -3, 7, 14, 21, and 28 dps after normalizing the score of each animal to its score at 2 dps.

**(b)** Same data as (a), showing raw mNSS results before normalization at -3, 2, 7, 14, 21, and 28 dps.

**(c)** Quantification of the percentage of ipsilateral hemisphere damage using MR images at 2, 7, 14, 21, and 28 dps of handled rats treated with a single or multidose of EVs after MCAO stroke.

**(d)** Same data as (c), showing total ischemic volume.

**(e)** Same data as (c), showing percentage of edematous expansion of the ipsilateral hemisphere.

All data was analyzed by two-way repeated measures ANOVA followed by the post hoc Holm-Sidak's multiple comparisons test (****$p<0.0001$). Bars show mean +/− SD. Each symbol represents a single rat: handled + EVs multi (*n=10*), handled + EVs single (*n=10*).
