## Supplementary material for "Recovery after human bone marrow mesenchymal stem cells (hBM-MSCs)-derived extracellular vesicles (EVs) treatment in post-MCAO rats requires handling associated with repeated behavioral testing": Uncropped images

### S1 RAW IMAGES

### Original uncropped blots from Fig 1b.

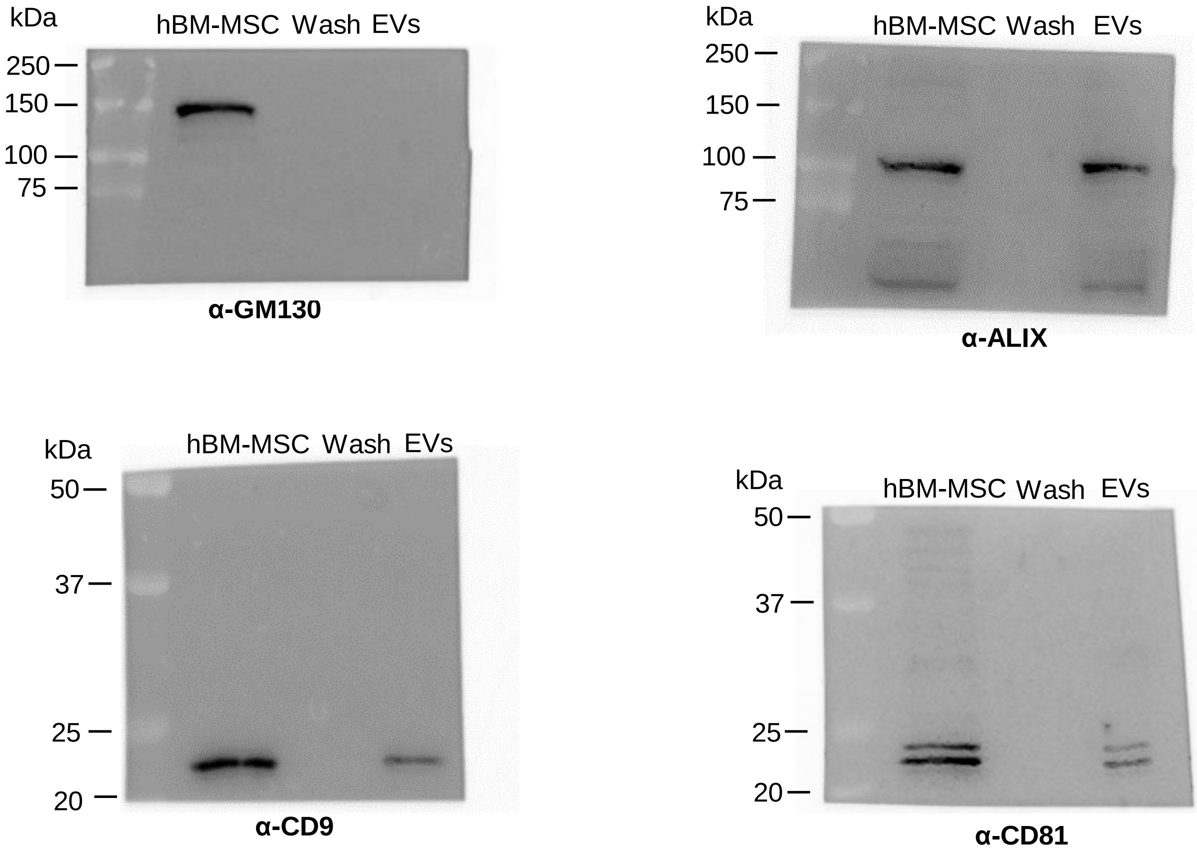


### Original uncropped TEM image from Fig 1c.

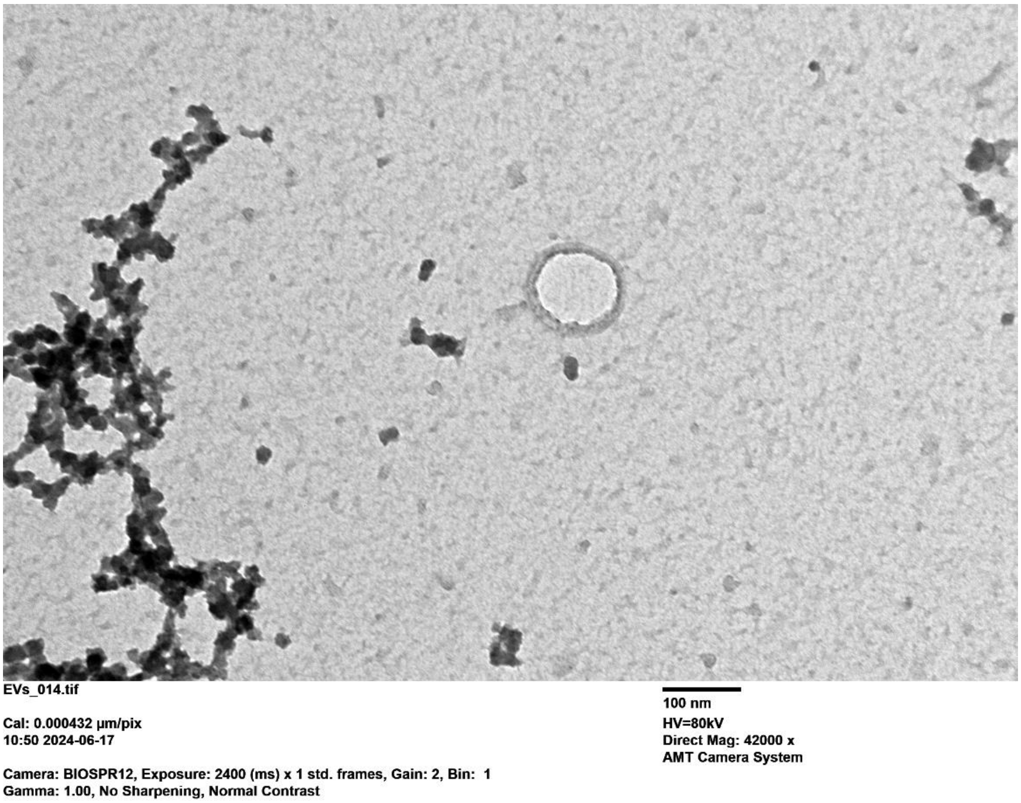
