## Supplementary tables (1-4) for "Recovery after human bone marrow mesenchymal stem cells (hBM-MSCs)-derived extracellular vesicles (EVs) treatment in post-MCAO rats requires handling associated with repeated behavioral testing"

**Table S1. Number of MCAO rats with treatments and behavioral assessments for each cohort.**

**Table S2. Description of all sensorimotor tests performed in this study.**

**Table S3. Description and score range of the independent tests in the modified Neurological Severity Score (mNSS).**

**Table S4. Description of possible scores (0 to 6) obtained during the beam balance test.**

**Table S1. Number of MCAO rats with treatments and behavioral assessments for each cohort.**

| Cohort | Intranasal treatment | Number of MCAO rats *(n)* | Behavior |
|---|---|---|---|
| **Non-handled** | Vehicle single dose | *n=12* | mNSS at -3, 2, 28, 56 dps |
| | EVs single dose | *n=13* | mNSS at -3, 2, 28, 56 dps |
| **Handled** | Vehicle single dose | *n=10* | mNSS, cylinder, corner, beam balance, grid walking, forelimb placement (vibrissae and proprioceptive) tests at -3, 2, 7, 14, 21, 28 dps |
| | EVs single dose | *n=10* | mNSS, cylinder, corner, beam balance, grid walking, forelimb placement (vibrissae and proprioceptive) tests at -3, 2, 7, 14, 21, 28 dps |
| **Handled** | Vehicle multidose | *n=10* | mNSS, cylinder, corner, beam balance, grid walking, forelimb placement (vibrissae and proprioceptive) tests at -3, 2, 7, 14, 21, 28 dps |
| | EVs multidose | *n=10* | mNSS, cylinder, corner, beam balance, grid walking, forelimb placement (vibrissae and proprioceptive) tests at -3, 2, 7, 14, 21, 28 dps |

**Table S2. Description of all the sensorimotor tests performed in this study.**

| Behavioral test | Description of the test |
|---|---|
| modified Neurological Severity Score (mNSS) | It evaluates sensorimotor activity and general neurological recovery after stroke |
| Beam balance test | It assesses fine motor coordination and balance |
| Corner test | It evaluates sensorimotor damage and postural asymmetry |
| Cylinder test | It measures spontaneous forelimb use to assess sensorimotor function and forelimb asymmetry |
| Grid walking test | It assesses fine motor function and limb coordination with accurate paw placement and grasping during locomotion |
| Proprioceptive forelimb placing test | It evaluates somatosensory and motor function with forelimb coordination, proprioception and tactile input |
| Vibrissae-evoked forelimb placing test | It evaluates somatosensory and motor function with unskilled reaching for a stable surface after unilateral vibrissae contact and visual trigger |

**Table S3. Description and score range of the independent tests in the modified Neurological Severity Score (mNSS).** A total score of 0 represents animals without deficits, while 16 points represent an animal with severe stroke symptoms. Only MCAO animals with striatal and cortical involvement and a mNSS equal to or above 9 were included in the experimental cohorts.

| **mNSS – Description for independent tests** | **Description of score** | **Score range** |
|---|---|---|
| **Postural signs** – *lift the rat by the tail in the air to check how it moves* | Symmetric forelimb extension when lifted by the tail<br>Forelimb flexion only<br>Forelimb flexion and thorax twisting | 0<br>1<br>2 |
| **Gait dysfunction** – *leave the animal walk-free on the table and observe how it walks* | Walking straight<br>Walking toward the contralateral side of the stroke<br>Alternate circling and walking straight<br>Alternate circling and walking toward the contralateral side<br>Circling and/or other gait disturbance | 0<br>1<br>2<br>3<br>4 |
| **Response to tail pull** – *grab the rat by the tail, leaving both front legs on the table. Pull up the back legs and observe how the rat moves* | Symmetric movement<br>Asymmetric movement (circling) | 0<br>1 |
| **Proprioceptive forelimb placing** – *check how the contralateral front paw is placed on the table when the animal walks* | Normal placing<br>Weak or delayed (<2s) placing of contralateral forelimb<br>Forelimb hanging | 0<br>1<br>2 |
| **Resistance to lateral displacement** – *both hands close to the animal, move the rat to the right or the left and check its resistance* | Normal symmetric resistance<br>Weakened resistance on the paretic side<br>No resistance on the paretic side | 0<br>1<br>2 |
| **Wire grasping strength** – *grab the animal by the tail and let it grab a wire with its front legs. Check its strength by grabbing the wire while pulling its tails* | Symmetric power<br>Asymmetric power | 0<br>1 |
| **Grasping reflex** – *grab the animal by the tail and let it grab a wire with its front legs. Check its forelimb reflex while grabbing the wire* | Grasps stick when palmar forepaw gently touched<br>No grasping | 0<br>1 |
| **Spontaneous activity** – *leave the animal walk-free on the table and observe how its exploratory activity* | Normal or near-normal exploratory and grooming behavior<br>Reduced locomotion and spontaneous limb movements<br>Responsive to stimuli only (tactile, auditory) - pretty lethargic<br>Immobile and unresponsive to stimuli and/or absent acoustic startle reflex - pretty lethargic | 0<br>1<br>2<br>3 |
| | **Maximum score** | 16 |

**Table S4. Description of possible scores (0 to 6) obtained during the beam balance test.** A score of 0 represents animals without deficits, while a score of 6 represents rats with severe stroke symptoms.

| Beam balance test – Description of score | Score |
|---|---|
| Animal balances with steady posture | 0 |
| Animal grasps side of the beam | 1 |
| Animal hugs the beam and one limb falls down from the beam | 2 |
| Rat hugs the beam and two limbs fall down from the beam, or spins on beam (>60 sec) | 3 |
| Animal attempts to balance on the beam but falls off (>40 sec) | 4 |
| Animal attempts to balance on the beam but falls off (>20 sec) | 5 |
| Animal falls off: no attempt to balance or hang on the beam (<20 sec) | 6 |
